# Disease persistence on temporal contact networks accounting for heterogeneous infectious periods

**DOI:** 10.1101/401158

**Authors:** Alexandre Darbon, Davide Colombi, Eugenio Valdano, Lara Savini, Armando Giovannini, Vittoria Colizza

## Abstract

The infectious period of a transmissible disease is a key factor for disease spread and persistence. Epidemic models on networks typically assume an identical average infectious period for all individuals, thus allowing an analytical treatment. This simplifying assumption is however often unrealistic, as hosts may have different infectious periods, due, for instance, to individual host-pathogen interactions or inhomogeneous access to treatment. While previous work accounted for this heterogeneity in static networks, a full theoretical understanding of the interplay of varying infectious periods and time-evolving contacts is still missing. Here we consider an SIS epidemic on a temporal network with host-specific average infectious periods, and develop an analytical framework to estimate the epidemic threshold, i.e. the critical transmissibility for disease spread in the host population. Integrating contact data for transmission with outbreak data and epidemiological estimates, we apply our framework to three real-world case studies exploring different epidemic contexts – the persistence of bovine tuberculosis in southern Italy, the spread of nosocomial infections in a hospital, and the diffusion of pandemic influenza in a school. We find that the homogeneous parameterization may cause important biases in the assessment of the epidemic risk of the host population. Our approach is also able to identify groups of hosts mostly responsible for disease diffusion who may be targeted for prevention and control, aiding public health interventions.

## 1 Introduction

Mathematical modeling of infectious diseases provides an important tool in understanding patterns and determinants of disease spread [1, 2]. The foundations of this field rest upon *compartmental models*, in which the population is divided in different classes (compartments) according to the individual health status and disease progression is modeled through transitions between compartments. This allows the mathematical description of the epidemic outcome, using, for instance, coupled differential equations [2, 3]. While usually being tractable, this framework, however, often relies on two important simplifying assumptions. The first is the homogeneous mixing approximation for which the probability of having a contact with an infectious host is identical for each individual of the population, neglecting any underlying social or spatial structure. Such approximation, unrealistic in many circumstances [4–9], was overcome by accounting for host contact heterogeneities in different ways. In this work we focus on the network epidemiology approach where each individual is represented by a node in a network and edges encode the interactions between nodes. The contact network can be static [10–14] or dynamic [7–9, 15–21], representing interactions that evolve in time. In both cases, network structure strongly influences the spreading process, since it depends on the interplay between the infection dynamics and the structural and temporal features of the networks.

The second assumption involves the recovery process, when infected hosts recover from infection either going back to the susceptible state, or acquiring immunity. Typically, models assume that all the infected individuals have identical constant recovery rate [2, 3]. This implies that the average duration of the infectious period (the interval during which an infectious host can transmit the disease) is the same for each individual and is exponentially distributed. Recovery is therefore described by an identical Markovian process for all individuals in the population that, while being analytically more tractable, can be unrealistic in many contexts. Individuals may indeed be characterized by different genetic [22] and immunogenetic [23] profiles, or may differ regarding age, medical treatment and vaccination [24]. These features, combined with specific epidemiological characteristics, may cause large deviations from the exponential distribution [25–27], and lead to large heterogeneities in the duration of the infectious period at the individual level. An increasing body of literature started to account for individual heterogeneities, considering mainly individual variations in infectivity [5, 28] and susceptibility [29–31]. Preliminary works incorporated this feature only in homogeneously mixed populations through integro-differential equations [10, 32], partial differential equations or through the method of stages [27, 33–37]. Only recently the effects of infectious period heterogeneity are analyzed on static networks, using message passing approaches [38, 39] and heterogeneous mean-field models [40]. These works provide a description of the Susceptible-Infectious-Recovered (SIR) model on static networks and the computation of the basic reproductive number *R*_0_ [2, 3]. These approaches however neglect the temporal dynamics of the network of contacts between hosts, which becomes particularly relevant if the time scale of the evolution of contacts is comparable to the one of the spreading process [7–9, 17, 18].

The aim of this work is to understand how heterogeneous durations of the infectious period impact the conditions for the spreading of an epidemic on a time-evolving contact network, in the particularly interesting scenario of the disease time scale being comparable to the one of the underlying network. By extending the infection propagator approach developed in [41, 42], we build an analytical framework that allows us to go beyond the two aforementioned assumptions providing an analytical form for the epidemic threshold, i.e., the critical transmission probability below which a pathogen would go extinct in the population. The epidemic threshold is a key epidemiological quantity as it can measure the vulnerability of a system to the introduction of a specific pathogen.

Lastly, we apply this methodology to evaluate the vulnerability of three different real-world systems: bovine tuberculosis in Southern Italy, nosocomial carriage of pathogenic bacteria in hospital facilities and pandemic influenza in closed settings. We show that using compartmental models with homogeneous infectious periods may introduce important biases in the estimate of the epidemic threshold and thus of the epidemic risk of a host population.

## 2 Methods

### Infection propagator for heterogeneous infectious periods

In this section, we introduce the analytical framework that allows us to compute the epidemic threshold for a spreading process on a time-varying contact network, considering that infected hosts have a constant rate of recovery, but each host is associated to an individual average infectious period.

We start from a Susceptible-Infectious-Susceptible (SIS) disease progression [2, 3], and assume a discrete time evolution for both the network and the spreading process. Each node of the network represents a host, and each link a time-resolved contact that is relevant to pathogen transmission. The temporal network is characterized by a finite number of snapshots *T* (network period) and formally it can be represented by a list of adjacency matrices whose element 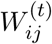 corresponds to the weight of the link made from node *i* to node *j* at time *t*. At each time step, infected nodes transmit the infection to the susceptible neighbors with a probability that depends both on the weight of the link and on the specific disease transmissibility *λ*.

The weight may correspond to different quantities (e.g. duration or strength of a contact) depending on the context under study, as we will see in the real-world systems addressed in this work. Infectious nodes can spontaneously recover with probability *μ*. Recovery is a Poisson process with an average infectious period *τ* = 1*/μ* that in the classic compartmental model formulation is the same for each individual.

Following the approach introduced in [41], the critical behavior of the epidemic spreading process is completely described by the following quantity, called infection propagator:

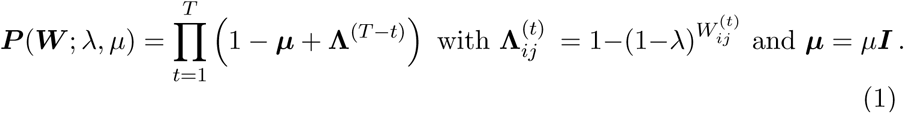

The sufficient and necessary condition for the existence of the asymptomatically stable disease-free solution is that the spectral radius of ***P*** (i.e. its lead eigenvalue) is smaller than 1. Therefore, for a given value of *μ*, the critical value of the transmissibility *λ*_*c*_ for which the spectral radius is equal to 1 corresponds to the epidemic threshold.

To account for hosts with individual average infectious period *τ*_*i*_, with *I* corresponding to the node index, we define 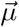 as a vector where each element *μ*_*i*_ is the inverse of the individual infectious period *μ*_*i*_ = 1*/τ*_*i*_. The infection propagator can then be written as:

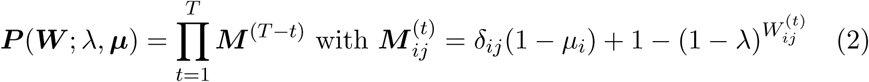

and the epidemic threshold expression becomes:

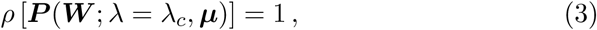

where ***μ*** is a diagonal matrix with 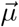 on the diagonal.

### Epidemic threshold computation

We compute the spectral radius of the infection propagator ***P***, given by the expression of Eq. (2) using the power iteration method. To compute the threshold, we find the zero for 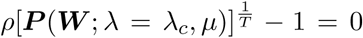 using Brent algorithm [43] where *λ* is a variable and *μ* is informed from epidemiological data. This procedure returns the value of *λ* for which the spectral radius is equal to one. A Python library to compute the critical transmissibility is publicly available online for interested researchers [44].

### Epidemic risk assessment in the heterogeneous vs. homogeneous cases

We compute the epidemic threshold with the infection propagator approach to assess the epidemic risk of a host population in three real case studies. In each one of them, actual epidemiological data allow us to inform the infection propagator ***P*** (***W***; *λ*, ***μ***) with empirical estimates for *μ*_*i*_, corresponding to the actual heterogeneous context. We then compare our findings with the epidemic threshold obtained under the assumption that all hosts have the same infectious period (referred to as the homogeneous case) obtained as a population average, i.e. τ_*all*_ = Σ_*i*_τ_*i*_/*N*, where *N* is the total number of hosts (nodes) in the population. This quantity enables a parameterization for the comparison of the two cases.

### Network properties

We define the activity potential of a node in an empirical network as the fraction of the number of time steps during which it makes contact with other nodes over the period of the network *T*, as defined in Ref. [21].

Other quantities used in this study are basic network measures, including the degree, i.e. the number of neighbors a given node has at time *t*, and the strength, i.e. the total sum of the weights on the connections that a given node establishes at time *t*. These quantities are also considered aggregated on a given time interval [45].

### Hellinger distance

To compare two distributions of the epidemic threshold obtained by varying underlying conditions in the host population under study, we use the Hellinger distance[46] that is defined for discrete distributions as follows:

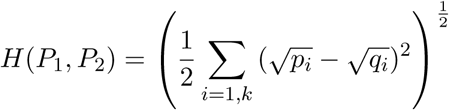

where *P*_1_ and *P*_2_ are discrete probability distributions over *k* discrete values: *P*_1_ = (*p*_1_*, …, p*_*k*_) and *P*_2_ = (*q*_1_*, …, q*_*k*_).

## 3 Results

### Bovine tuberculosis in Italy

Livestock infectious diseases are of primary interest for animal health and welfare and for the economy of a country, with the potential to produce devastating consequences [47]. Bovine tuberculosis is one the most widespread zoonotic diseases all over the World [48] and it is very difficult to contain since it can rapidly spread while unnoticed, before the onset of clinical signs in the animals [49]. Bovine tuberculosis is a notifiable disease in Europe. In Italy, in particular, it has affected cattle population for decades and still circulates in southern regions [50]. Here we focus on Puglia region in the south of Italy where outbreak data are available and outbreak duration was shown to vary depending on the production type of the affected premises.

We built the temporal network starting from the dataset of cattle trade movements of the whole population of bovines in Puglia obtained from the Italian National Database for Animal Identification and Registration that is managed by the Istituto Zooprofilattico Sperimentale dell’Abruzzo e del Molise (IZSAM) on behalf of the Italian Ministry of Health [51, 52]. Each time-stamped cattle movement record provides the animal unique identifier and the identifiers of both origin and destination premises. Here we consider the daily records of bovine trades in Puglia, from January 1, 2006 to December 31, 2012. In this region, 5,430 animal holdings displaced 136,206 bovines through 44,272 trade movements in the time period under study. In the network representation, each node corresponds to a farm, and directed links represent the animal movements, weighted with the number of bovines moved [53, 54].

Bovine displacements tend to concentrate in the countryside around the three majors cities of the region: Foggia, Bari and Lecce (Figure 1a). Outbreak locations correspond to these three geographic clusters, and also appear to be grouped according to the production type of the affected premises (i.e. meat, dairy, or mixed production). We define the monthly-aggregated average degree of an animal holding as the average number of trading partners it has in a month. We compute this quantity by averaging on all premises and averaging only on the ones trading at least once a month (defined as active, including both sales and purchases). The evolution of these quantities over time shows the low density of the network (Figure 1b). Each month active premises represent between 8 and 14% of the total number of premises and their average degree is around 3, leading to a very low overall average degree. Further analysis of the Italian cattle trade network properties were performed in previous works [9, 55, 56].

**Figure 1:**
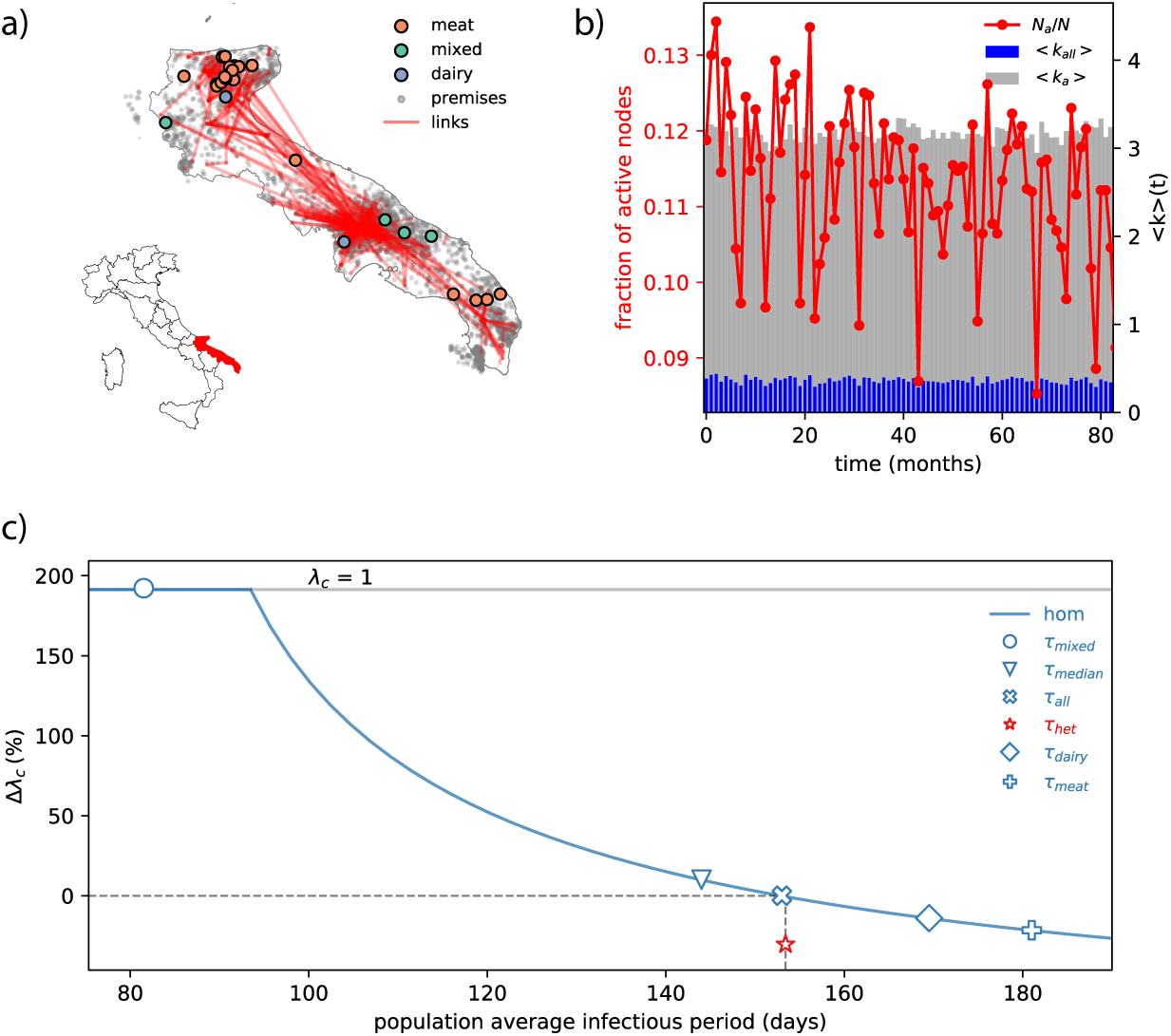
Bovine tuberculosis in Puglia, Italy. a) Aggregated network of the cattle trade movements in Puglia and location of bovine tuberculosis outbreaks in this region, in the period 1983 - 2015. Only the links with an aggregated strength higher than 30 are represented. Affected farms are identified according to their production type (dairy, meat, mixed). b) Activity patterns of the cattle trade network in time. The red line shows the proportion of active nodes. Blue bars show the monthly average degree of all nodes and grey bars show the monthly average degree of active nodes. c) Epidemic threshold estimate comparison between homogeneous and heterogeneous parameterizations. The plot shows the relative variation of the epidemic threshold (expressed in %) as a function of the population average infectious period. The reference for the relative variation is the epidemic threshold obtained in the homogeneous assumption obtained with the population average infectious period *τ*_*all*_ computed from the data (Table 1). The red star shows the epidemic threshold estimate in the heterogeneous case. Blue markers show particular cases of the threshold estimate in various homogeneous parameterizations. The blue line shows the full dependence of the epidemic threshold estimate in the homogeneous case as a function of the population average infectious period.

For the outbreak data we used records of bovine tuberculosis cases in cattle occurring in Puglia between 1983 and 2015. This data were collected by the Italian National Animal Health Managing Information System, an informatics system developed by IZSAM on behalf of the Ministry of Health [51, 52]. For each outbreak, we considered data reporting the identifiers of the affected animal holding, its production type, and the start and the end of the outbreak based on the date of first infection detection and date of clearing of the infection, respectively. As shown in Table 1, outbreak durations vary with production types. We split the premises population in groups corresponding to their production type, and we set to each node the data-driven infectious period corresponding to the associated production type.

**Table 1:** Estimated outbreak duration of bovine tuberculosis per production type of the affected farm.

| Production type | Outbreak duration (min-max) (days) [51, 52] |
| --- | --- |
| Meat | $\tau_{meat} = 181$ (74-2029) |
| Diary | $\tau_{dairy} = 170$ (65-274) |
| Mixed | $\tau_{mixed} = 82$ (41-292) |
| All | $\tau_{all} = 153$ |

Predicted epidemic risks in the homogeneous or heterogeneous case are rather different (Figure 1c). The epidemic threshold computed on heterogeneous durations of the outbreak durations is about 30% smaller than the estimate obtained with the population average. As a result, the homogeneous assumption overestimates the epidemic threshold. Even if we vary the population average infectious period (used in the homogeneous model), for example by assuming that all premises are either dairy or meat farms, we obtain a higher epidemic threshold, thus systematically underestimating the vulnerability of the livestock system. More in detail, we obtain the largest bias when we parameterize all premises as they were all of mixed production. Under this assumption, the obtained epidemic threshold is equal to 1, meaning that the pathogen would not be able to persist in the system, in clear contradiction with the current epidemiological status of Puglia with respect to bovine tuberculosis.

### Nosocomial infection carriage in hospital facilities

The definition of *nosocomial infection* applies both to infections acquired by patients while being in a healthcare facility, and to occupational infections of medical staff [57]. Their control is a great concern in public health. Several pathogens cause nosocomial infections, such as viruses (e.g. nosocomial influenza, rotaviruses), or pathogenic and nonpathogenic bacteria (e.g. *Mycobacterium Tuberculosis*, *Pseudomonas Aeruginosa*, *Klebsiella Pneumoniae*, *Enterococcus*, *Staphyloccocus Aureus*) [58]. While host colonization can last for months in the absence of intervention[59], it can be dramatically shortened by detection and intervention protocols, or by the hygiene measures adopted by hospital personnel [60]. In addition, patients are known to take longer to clear nasal colonization than healthcare workers [60, 61]. In order to understand how these differences may affect the vulnerability of a hospital setting to nosocomial infections, we focus here on the case of *Staphylococcus Aureus* (*S. Aureus*), for which colonization durations are documented [60–62].

Close proximity interactions are acknowledged to be a route along which nosocomial infections may be transmitted [61, 62]. Here we use a temporal network of contacts within a hospital ward in Lyon, France, collected by the SocioPatterns project [63, 64]. The dataset records face-to-face interactions among patients and hospital workers over 4 days, at a 20 second resolution. It includes also individual information that allow us to divide the population into four groups: patients, nurses, medical doctors and administrative personnel (see Table 2). Spatial constraints appear to be major drivers in network topology, as, for instance, patients are mostly confined to rooms, and administrative personnel to offices. Nurses, moreover, are the most connected, sharing links with all other groups and among themselves (Figure 2a). The number of contacts follows a daily cycle, featuring peaks during daytime, and troughs at night (Figure 2b).

**Table 2:** Number and proportion of individuals in each class in the hospital network and corresponding estimated carriage duration for *S. Aureus*.

| Role | #nodes (%) | Carriage duration (days) [60–62] |
| --- | --- | --- |
| Patients | 29 (38.67) | $\tau_p = 10$ |
| Nurses | 27 (36) | $\tau_n = 2.5$ |
| Medical Doctors | 11 (14.66) | $\tau_m = 2.5$ |
| Administrative personnel | 8 (10.67) | $\tau_a = 2.5$ |
| All | 75 (100) | $\tau_{all} = 5.4$ |

**Figure 2:**
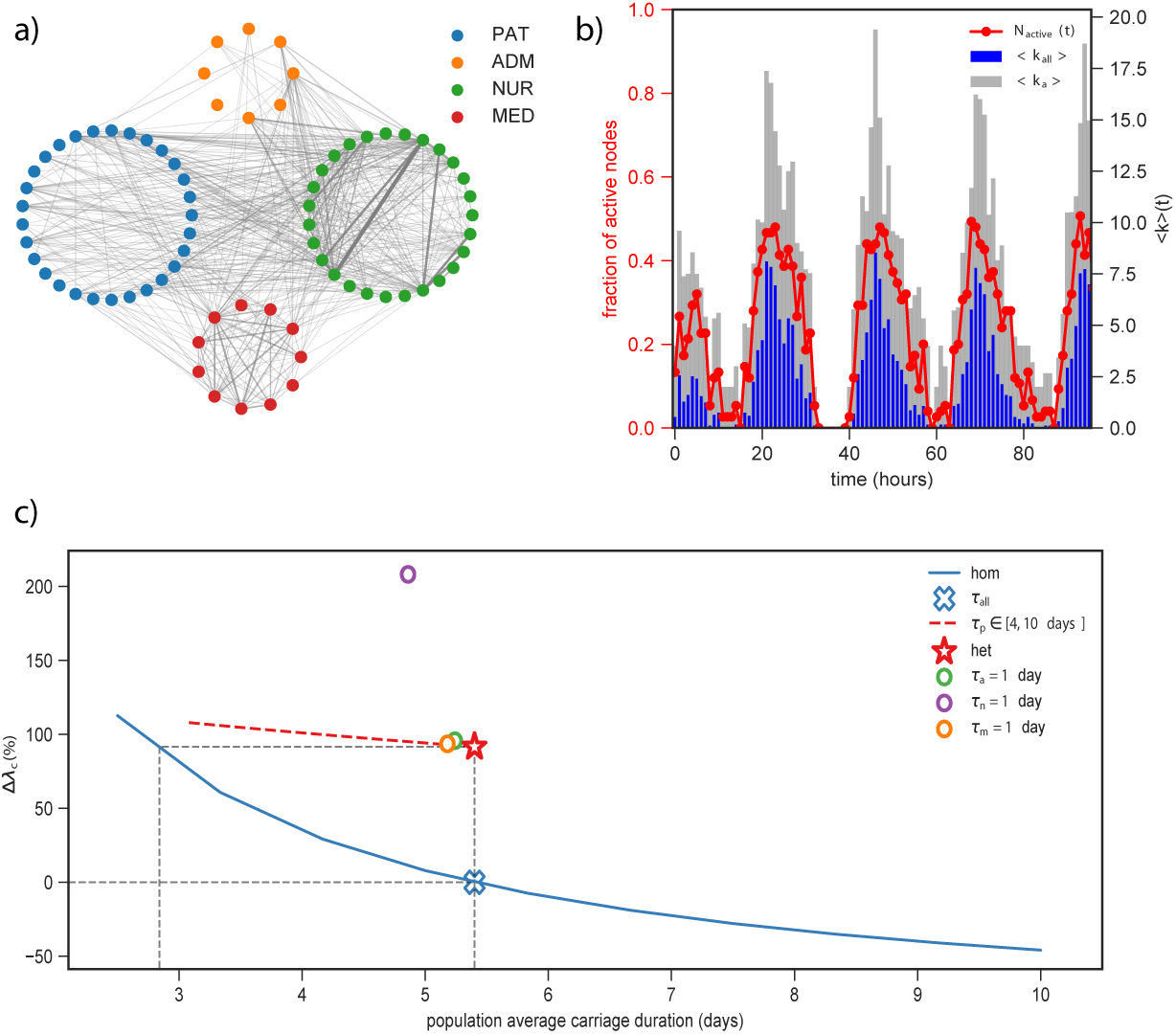
*S. Aureus* diffusion in a hospital ward. a) Aggregated network of proximity contact in the hospital ward. Nodes are grouped by class of individuals and links’ width and color are proportional to links’ weight, measuring the total amount of time associated with each contact. b) Activity patterns of the hospital network in time. The red line shows the proportion of active nodes. Blue bars show the hourly average degree of all nodes and gray bars show the hourly average degree of active nodes. c) Epidemic threshold estimate comparison between homogeneous and heterogeneous parameterizations. The plot shows the relative variation of the epidemic threshold (expressed in %) as a function of the population average infectious period. The reference for the relative variation is the epidemic threshold obtained in the homogeneous assumption (blue cross) obtained with the population average infectious period *τ*_*all*_ computed from the data (Table 2). The red star shows the epidemic threshold estimate in the heterogeneous case. Circles correspond to the estimates obtained when reducing the carriage duration of each class of hospital personnel to 1 day, in the heterogeneous parameterization. The red dashed line show the estimates obtained when reducing the carriage duration of patients in the range of 4 to 10 days, in the heterogeneous parameterization. The blue line shows the full dependence of the epidemic threshold estimate in the homogeneous case as a function of the population average infectious period.

Analogously to the case of bovine tuberculosis, we compare the epidemic threshold obtained under homogeneous and heterogeneous parameterizations of the infection propagator approach. In the heterogeneous case, patients and personnel colonization durations are set to respectively 10 and 2.5 days[60–62] while *τ*_*all*_ = 5.4 days (Table 2). The epidemic risk analysis presented in Figure 2c shows that the epidemic threshold in the heterogeneous case yields a 100% relative variation with respect to the homogeneous parameterization with *τ*_*all*_. This hints at the presence of a group of potential hosts that are critical to the spread of the pathogen, but have a short infectious period. To obtain the same estimates of the epidemic threshold of the heterogeneous case with the homogeneous parameterization we would need to reduce the average infectious period to *τ*_*all*_ =2.8 days.

To understand the contribution of each class of individuals to the epidemic risk, we computed the epidemic threshold in the heterogeneous parameterization by reducing the carriage duration parameter of a single class at a time for each of the four classes. For the hospital staff, we set the carriage duration to 1 day, and for the patients, we explored a reduction in their carriage duration from 10 to 4 days. This allows us to understand the performance of an intervention aimed at reducing the epidemic risk for the ward, using, for instance, more frequent hand washing for a given class of staff, or enhanced screening and treatment of patients. We observe that while the intervention on administrative personnel or medical doctors would have almost no impact on the epidemic threshold, targeting nurses would dramatically increase the epidemic threshold, yielding a relative variation larger than 200% with respect to the homogeneous assumption. This highlights the important role of nurses in the network of contacts, and their potential of largely facilitating the dispersal of nosocomial infections in the hospital setting [62, 65–67].

### Pandemic influenza in closed settings

Influenza is a respiratory infectious disease that spreads through proximity contacts, and affects 10% to 30% of European population every year [68]. High-risk individuals, such as elderly or immune-deficient, may experience a severe form requiring hospitalization. Particular attention is given to schoolchildren [69, 70] since epidemiological evidence suggests that they are critical in the early transmission chains of the disease favoring then the diffusion in the general population. Hosts infected with influenza may develop symptoms or may be asymptomatic, also exhibiting a shorter infectious period in absence of symptoms[24]. We aim at understanding how the presence of asymptomatic individuals impacts the pandemic risk for a population of students at a school.

We build the temporal network from data reporting time-resolved contacts in a French school [71, 72] collected by the SocioPatterns project [63], over a period of 2 days, at a resolution of 20 seconds. As for hospital facilities, proximity sensors were used to record face-to-face contacts between individuals. The resulting network is composed of *N* =242 individuals divided into 11 classes, corresponding to ten classes distributed on five consecutive grades, plus teachers. The network exhibits strong community structure, as children connect more within the same class or the same grade (see Figure 3a). While teachers connect with students from different classes, those links have a lower weight, as they occur less frequently and are shorter in time. The hourly activity timeline shows clearly the three main breaks of the day (Figure 3b). The proportion of active nodes is very high, showing a high degree of interaction among students [73, 74]. It is possible to distinguish the morning and afternoon breaks, characterized by a proportion of active nodes close to 1, from the lunch break where students have lunch or leave the school to eat at home decreasing the network activity. Detailed analysis of the network were reported in previous work [71, 72].

**Figure 3:**
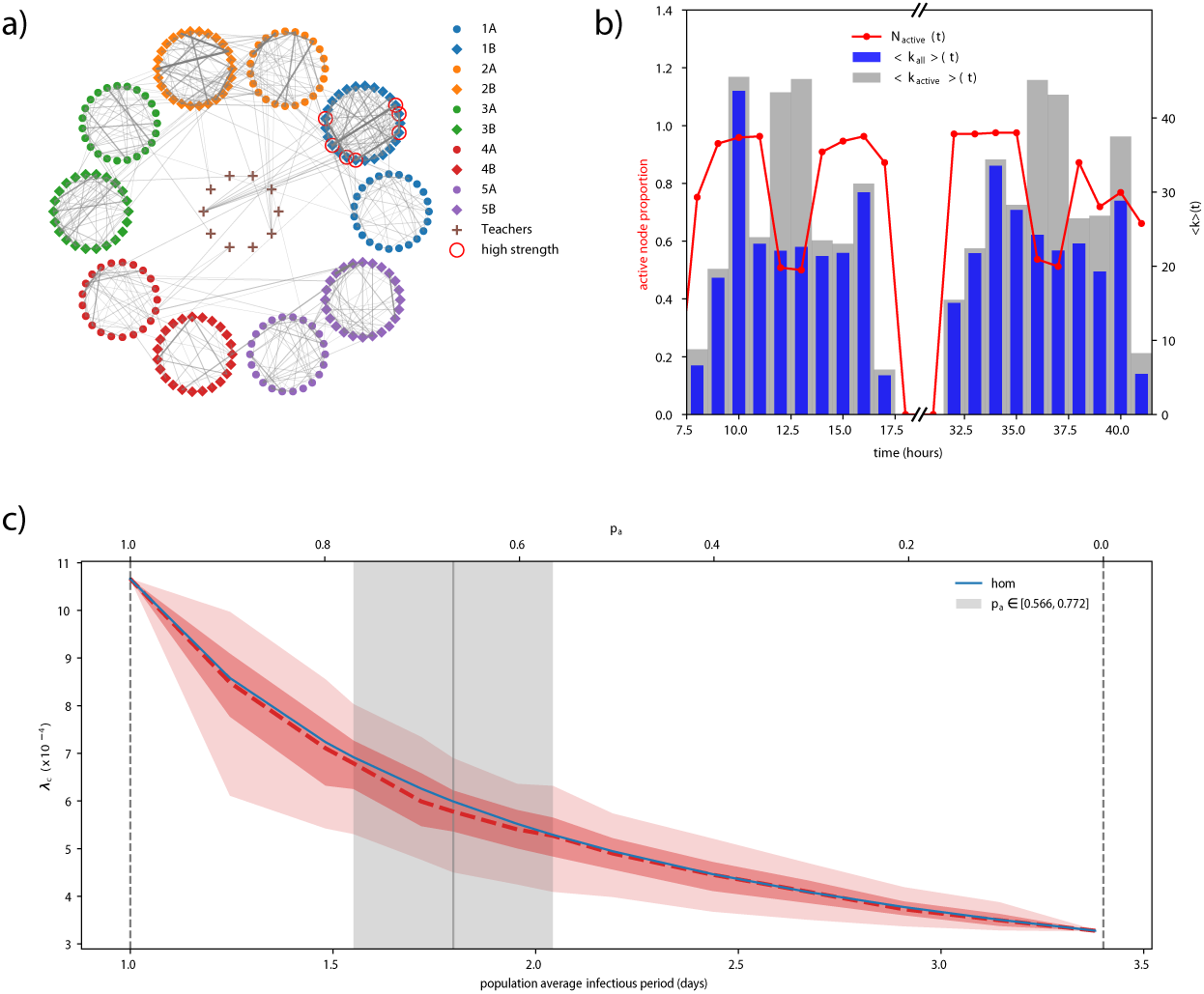
Pandemic influenza in a school setting. a) Aggregated network of proximity contacts in the school. Nodes are grouped by class of individuals and links’ width and color are proportional to links’ weight, measuring the aggregated amount of time associated with each contact. b) Activity patterns of the school network in time. The red line shows the proportion of active nodes. Blue bars show the hourly average degree of all nodes and gray bars show the hourly average degree of active nodes. c) Epidemic threshold estimate comparison between homogeneous and heterogeneous parameterizations. The plot show the epidemic threshold as a function of the population average infectious period, for both homogeneous (blue) and heterogeneous (red) parameterizations. The red shaded areas show the 95% (lighter red) and 50% (darker red) confidence intervals for the epidemic threshold estimate obtained when randomizing on nodes that are symptomatic or asymptomatic in the heterogeneous case. The gray vertical area indicate the range of population average infectious period corresponding to *p*_*a*_ varying in the empirically estimated 95% confidence interval.

We model the epidemic using infectious period estimates from 2009 influenza pandemic [24, 75], yielding 3.4 days for symptomatic infections and 1 day for asymptomatic infections (Table 3). We investigate proportions of asymptomatically infected schoolchildren (*p*_*a*_) in agreement with empirical estimates, i.e. *p*_*a*_ = 0.669 (95% CI: 0.556-0.772) [76]. In addition to these values, we also explore the full range from *p*_*a*_ = 0 to *p*_*a*_ = 1 for a comprehensive analysis. For a given proportion *p*_*a*_ of asymptomatic individuals we compute the epidemic threshold randomly extracting the asymptomatic nodes *p*_*a*_*N* in the population. Repeating this operation 500 times, we obtain a distribution of epidemic threshold values for each value or *p*_*a*_ depending on the assumed symptomatic response of the host population of schoolchildren.

**Table 3:** Estimated infectious period per symptomatic or asymptomatic influenza infections. The population average *τ*_*all*_ depends on the assumed proportion *p*_*a*_ of asymptomatic individuals in the schoolchildren population.

| Type of infection | Infectious period (days) [24] |
| --- | --- |
| Symptomatic | $\tau_s = 3.4$ |
| Asymptomatic | $\tau_a = 1$ |
| All | $\tau_{all} = \tau_{all}(p_a)$ |

Figure 3c reports the results of the comparison between the homogeneous and heterogeneous cases. We observe that without any asymptomatic individual (i.e. *p*_*a*_ = 0 and population average infectious period equal to 3.4 days) the threshold is lower than when considering the presence of asymptomatic, as expected. While it may be natural to think that asymptomatic individuals may create unnoticed paths of infection, their shorter infectious periods indeed plays a role in reducing the epidemic vulnerability of the population. When we consider the heterogeneous parameterization, we find that the homogeneous epidemic threshold is almost equal to the median heterogeneous one. This observation is not only true for values of *p*_*a*_ in the confidence interval of the empirical estimates, but also for the whole range from 0 to 1. There are, however, significant fluctuations around the median value, whose width corresponds to approximately 40% of the homogeneous estimate. The epidemic risk assessment assuming homogeneous infectious periods in the population may therefore introduce a large bias leading either to underestimating or overestimating population’s risk to infection.

We now want to understand the mechanisms responsible for those fluctuations, specifically, by identifying the hosts that play a key role in shaping the vulnerability. In what follows, we fix the proportion of asymptomatic individuals to the empirical estimate *p*_*a*_ = 0.669 [76]. For each node, we determine the two threshold distributions obtained considering the given node as either symptomatically or asymptomatically infected. We then compute the Hellinger distance between these two distributions to quantifies their dissimilarity: the higher the distance is, the more different the two distributions are (see Methods). This quantity will be higher for nodes whose status has a larger impact on the epidemic threshold. The distribution of the Hellinger distance of all nodes, shown in Figure 4a, is bimodal and the subset of nodes mainly responsible for the epidemic threshold variation is clearly visible. It is a small group composed of only 7 nodes, corresponding to 2.9% of the school population. In Figure 4b we show two examples of the distributions obtained for a node selected in each mode of the distance distribution.

**Figure 4:**
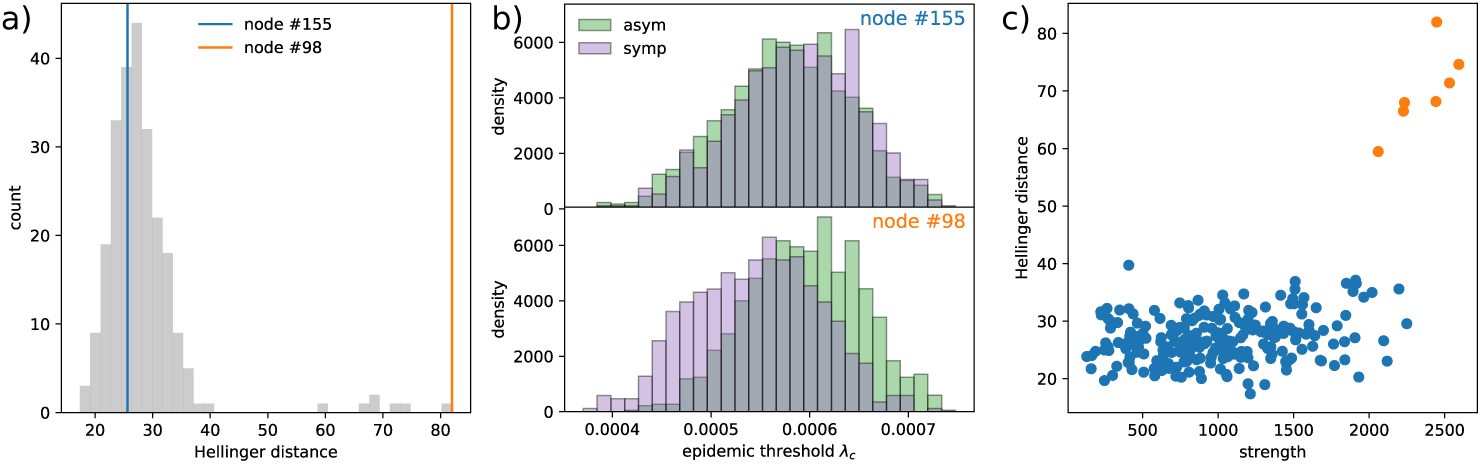
Identification of nodes mostly contributing to the large fluctuations obtained with the heterogeneous parameterization compared to the homogeneous case. a) Distribution of the Hellinger distance computed on the epidemic threshold distributions obtained by considering one at a time each node in the symptomatic or asymptomatic classes. The blue and the orange lines show examples of distance values obtained for two nodes, one close to the peak of the distribution (blue) and the other being an outlier (orange). b) Epidemic threshold distributions depending on symptomatic status following infection (purple for symptomatic infection, green for asymptomatic infection), for the two nodes highlighted in panel (a). c) Hellinger distance as a function of the node strength. Orange points correspond to nodes with outlier Hellinger distance values.

We also observe that these 7 nodes belong to the same class (see circled nodes in Figure 3a) and do not correspond to hubs in the whole network. They are likely to be a single group of friends. In order to highlight the patterns making these specific nodes more responsible than others for the threshold variation, we investigate various network properties. We find that these nodes exhibit higher strength (Figure 4c) and activity potential (not shown).

## 4 Discussion

Mathematical models have been highly successful in the study and understanding of infectious disease epidemics [2, 3]. Some simplifying assumptions, however, have sometimes hindered their applicability to real world scenarios. Our work helps adding realism to modeling a wide class of highly relevant diseases, as it provides a synthetic and solvable framework for considering individual hosts (or host classes) that have different characteristic infectious periods. By building on the methodology described in [41, 42], we have defined and computed the epidemic threshold with arbitrarily heterogeneous recovery probabilities, assuming them to be constant. We then applied our analytical framework to three relevant case studies, showing that we can successfully measure how such heterogeneity impacts the vulnerability of a particular population to disease introduction, considering bovine tuberculosis in southern Italy, nosocomial infections in hospital wards, and influenza-like epidemics in schools.

We stress that, while an increasing body of literature has already demonstrated the importance of including contact structure [4–16] and host-specific heterogeneities [25–31, 37, 77], few studies have considered the combined impact of these two factors in the estimation of the epidemic risk. Theoretical works that include host-specific heterogeneities were previously carried while assuming homogeneously mixed population [10, 27, 32, 33, 37] or using a static contact network [38–40], neglecting the possible interplay between the spreading process timescale and the contact evolution.

Our findings suggest that obtaining an accurate, individual or class-specific, estimate of the infectious period is an important step for building realistic spreading models. Indeed we have shown that if we set the same recovery probability to all hosts, we may greatly bias the estimate of the vulnerability, with the sign of the error depending on the specific setting. For bovine tuberculosis in Puglia, any homogeneous parameterization of the outbreak duration predicts a lower epidemic risk with respect to the heterogeneous case. In particular, when the shortest recorded outbreak duration is considered in the homogeneous approximation (*τ* = *τ*_*mixed*_), the disease is predicted not to be able to persist in the system, in evident contrast with the documented endemic presence of bovine tuberculosis in the region. On the contrary, in the case of the *S. Aureus* infection within a hospital, the classic homogeneous assumption causes to substantially overestimate the epidemic risk, with respect to the heterogeneous case. The presence of medical personnel with a shorter carriage duration is able to counteract the spreading potential of patients assumed to be carrying the pathogen for a longer time, and to further reduce the risk with respect to a homogeneous parameterization with the population average. Finally, for pandemic influenza in school settings, even though the median epidemic threshold obtained from the heterogeneous assumption matches the homogeneous result, we observe important fluctuations that may lead to either underestimating or overestimating vulnerability.

All these findings highlight the importance of properly accounting for heterogeneous infectious periods, as ignoring this feature may lead to a biased estimate of the epidemic risk, and consequently inefficient control strategies [78].

In addition, despite the fact that the responsibility of more active nodes in the epidemic risk has already been proven [45, 56], our approach was able to highlight them and show their contribution to population vulnerability, depending on their individual infectious period and their network properties. This can have clear implications in devising targeted interventions. For example, we have shown that in the hospital ward hygiene practices that reduce carriage duration in nurses can lead to a strong reduction in nosocomial risk. No impact is instead observed reducing the carriage duration of the administrative personnel or medical doctors. These findings are explained by the highly connected role of nurses within the hospital facility, given they interact considerably with both patients and other hospital personnel. As such, they represent an ideal target group for prevention measures aimed at lowering the risk for pathogen diffusion [64, 79–81]. Also in the case of pandemic influenza in school our approach allows the identification of those individuals who mostly contribute to disease circulation and persistence.

Our findings provide useful insights for the understanding of host heterogeneities in disease spread and can be used to build more realistic data-driven mathematical approaches for real case scenarios and targeted control measures. However, several key theoretical and practical issues are still to be addressed. First, empirical evidence shows that asymptomatically infected individuals with influenza tend to be less infectious than those who develop symptoms [24]. Our approach, however, assumes that disease transmissibility would not vary depending on host symptoms status, as our study was restricted to the role of heterogeneity of infectious periods only. Such variation can be taken into account by introducing a variation of the weights of the contacts established by an asymptomatic infectious individual while infected. The framework with varying transmissibility may also apply to other epidemic contexts, such as e.g. the spread of nosocomial infections. Nurses and doctors indeed are expected to adopt hygiene measures (e.g. use of disposable gloves, hand washing, etc.) that can reduce their transmissibility if infected.

Second, our study considered a basic intervention expressed with the reduction of the carriage duration for medical personnel in the hospital. More realistic interventions may however be considered, as e.g. pharmaceutical treatments aimed not only at reducing the infectious period but also the probability of transmission of the disease (e.g. antiviral treatments for influenza infections [24], or the use of antibiotics for bacterial infections). In addition, recent work highlighted the role of heterogeneous carriage duration of bacterial strains in ruling the relative abundance of each strain under antibiotic treatments [82]. Accounting for these aspects would allow a more realistic design of targeted interventions aiming at raising the epidemic threshold in the most efficient way.

Finally, more realistic and pathogen-specific distributions of infectious periods should be further considered, differently from the exponential distributions with heterogeneous individual infectious periods assumed here. This would likely require a radical re-design of the infection propagator approach to address the inclusion of such distributions. These various future directions would help the theoretical understanding of disease spreading processes with increasingly realistic epidemic models.

